# Isotope-based visualization of element distribution in phloem provides functional evidence for the operation of SOS1 Na^+^/H^+^ exchangers in mature zones of Arabidopsis root

**DOI:** 10.1101/2022.04.29.489905

**Authors:** Ryohei Sugita, Takaaki Ogura, Natsuko I. Kobayashi, Muhammad B. Gill, Lana Shabala, Tomoko M. Nakanishi, Sergey Shabala, Keitaro Tanoi

## Abstract

In the phloem, various solutes such as photosynthates, mineral nutrients, or toxic elements move from the source organs to sinks. One of them is the root. While this process may mediate significant quantities of solutes delivered to the root, it is unclear how the solutes are distributed along the root axis. To elucidate the distributing pattern of solutes, we visually analyzed the movement of solutes in Arabidopsis roots using a radioisotope imaging technique. The distribution patterns fell into four different groups: (1) root tip accumulation for ^14^C-photosynthates, ^28^Mg^2+^, ^32^PO_4_^3-^, and ^35^SO_4_^2-^; (2) homogenous distribution along the root axis for ^42^K^+^ and ^137^Cs^+^; (3) no detectable accumulation in the root for ^45^Ca^2+^ and ^59^Fe^2+^; and (4) transient accumulation in the root for ^22^Na^+^. The latter phenomenon was explored in more detail using Arabidopsis knockout mutants lacking functional sodium efflux transporter SOS1 (operating as Na^+^/H^+^ exchanger). By utilizing a non-invasive microelectrode MIFE ion flux measuring technique, we found that Na^+^ efflux was active in the mature root zone of wild-type Arabidopsis plants but not in root apex as initially thought, and that in *sos1* mutants lacking functional Na^+^/H^+^ exchangers, shoot-derived ^22^Na^+^ remained in the root. These findings challenge the notion that Na^+^ exclusion via SOS1 is confined to the root apex and demonstrates the power of combining the radioisotope imaging technique and the MIFE ion flux measuring technique to study the kinetics of ion transport in the root and root-to-shoot communication.

## Introduction

In plants, photosynthates and minerals in source organs are efficiently translocated to developing organs via the phloem (Beevers, 1969; Muchow et al., 1976; Taiz and Zeiger, 2002). Approximately 30% of all carbohydrates produced by leaves go to roots (Hole and Dearman, 1990; Whiley et al., 1993; Heuvel et al., 2002; Remus and Augustin, 2016), and up to 80% of K^+^ taken by roots and delivered to the shoot can be cycled back via the phloem (Marschnert et al., 1997). Various methods have been developed in studies of phloem transport, and those methods can be categorized by the object of analysis, such as determination of the solute composition in the phloem, determination of the mass flow rate in the phloem, and tracking of the solute movement via the phloem. To analyze the solute composition in the phloem, the phloem sap is collected from a cut end of a stem (Balachandran et al., 1997; Kehr et al., 1999; Fiehn, 2003; Tetyuk et al., 2013). Alternatively, phloem sap can be collected using an insect stylet (Kawabe et al., 1980; Fukumorita and Chino, 1982; Aoki et al., 2005), which has less chance of exudate contamination from other cells. With the collected phloem sap, changes in the chemical form of compounds due to the metabolism can be tracked by the stable isotope analysis (Hegeman et al., 2007; Tcherkez et al., 2011). The mass flow rate in the phloem has been analyzed with the heat pulse method, where multiple probes inserted into the plant measure the heat transfer between the probes (Ziegler and Vieweg 1961; Čermák et al. 2004). To track the movement of solutes via the phloem, tracer-based methods are often employed. The fluorescent dye tracer method, in combination with confocal laser microscopy, allows non-invasive real-time imaging of phloem mass flow with high resolution (Oparka et al., 1994; Write and Oparka, 1996). Compared to the fluorescent dye tracer, the radioactive tracer method has some advantages; tracers of the same chemical form as the traced compounds are available, and quantification of radioactive tracers is often feasible since radiations from most radioisotopes used in life science are strong enough to penetrate tissues of several millimeters thickness (Sugita et al., 2014). Two radioisotope imaging systems, <u>P</u>ositron <u>E</u>mitting <u>T</u>racer <u>I</u>maging <u>S</u>ystem (PETIS; Fujimaki et al., 2015) and <u>P</u>lant <u>T</u>omographic <u>I</u>maging <u>S</u>ystem (PlanTIS; Schmidt et al., 2020), are nondestructive radioisotope imaging systems that apply medical imaging technology to plant imaging. The principle of these imaging systems is based on the detection of two γ-rays originating from a positron, and the strong tissue penetration capacity of γ-rays allows the imaging of roots and root vegetables in soil (Jahnke et al., 2009). However, the use of these imaging systems is limited to radioisotopes that emit positrons only. Another system for plant imaging, the <u>R</u>eal-time <u>R</u>adioisotope <u>I</u>maging <u>S</u>ystem (RRIS), can detect β-, X-, and γ-rays since its principle of radiation detection is to convert radiations into light (Sugita et al., 2014). This enables the use of various radioisotopes, and the RRIS previously visualized the movement of various important solutes for plants after root uptake in Arabidopsis (Sugita et al., 2016). With this imaging system, the movement via the phloem can also be tracked for multiple types of solutes.

For most elements, mechanisms of phloem unloading and following transport in roots are yet to be elucidated. One of those elements is sodium (Na), which inhibits the activities of many cytosolic enzymes that depend on potassium (Greenway and Osmond, 1972; Shabala and Cuin, 2008; Wu et al., 2018). Phloem unloading of sodium ion (Na^+^) is a part of Na^+^ recirculation, which leads to Na^+^ exclusion from shoots (Blom-Zandstra et al., 1998). Recirculation of Na^+^ consists of phloem Na^+^ loading in shoots and unloading in roots, both of which are thought to be mediated by members of class I high-affinity K^+^ transporters (HKT1) (Berthomieu et al., 2003; Kobayashi et al., 2017; Venkataraman et al., 2021). In roots, Na^+^ unloaded from the phloem should be sequestrated into vacuoles or excluded back to the rhizosphere because Na^+^ accumulation in the cytosol could compromise root functions. For the sequestration, Na^+^ transport into vacuoles by tonoplast-based Na^+^/H^+^ exchanger NHX1 (Gaxiola et al., 1999; Apse et al., 1999) and Na^+^ retention in vacuoles by the efficient control of Na^+^ leak channels; slow-activating (SV) and fast-activating (FV) channels (Shabala et al., 2020) are considered relevant. For the exclusion, plasma membrane Na^+^/H^+^ antiporter SOS1 (Shi et al., 2002) is the only responsible transporter characterized to date. However, there is some apparent inconsistency regarding the expression pattern of *SOS1* in roots and the mode of their operation. Shi et al. (2002) reported *SOS1* gene expression specific to the pericycle and the root tip epidermis using transgenic Arabidopsis plants, while relatively ubiquitous expression in the root was detected by cell type-specific microarray expression profiling (Brady et al., 2007). Despite this controversy, the dominant view is that SOS1-mediated Na^+^ exclusion operates predominantly in the root apex (Munns and Tester 2008; van Zelm et al. 2020; Zhao et al. 2020). This notion, however, is counterintuitive, as the root apex represents only a small fraction of the bulk of the root. So, what is the point for the plant to transport Na^+^ along the main axis to be excluded in some confined space?

In the current study, we utilized the RRIS to visually analyze the phloem unloading of multiple solutes in roots of intact Arabidopsis seedlings and revealed four distinct phloem unloading patterns for the analyzed solutes. For ^22^Na transport, transient accumulation of shoot-derived ^22^Na^+^ was observed in wild-type Arabidopsis roots, while in *sos1* mutants lacking functional Na^+^/H^+^ exchangers, shoot-derived ^22^Na^+^ remained in the root and was not expelled. Intriguingly, this Na^+^ exclusion in wild-type roots was observed in the mature root zone but not in the root apex, challenging the classical view and providing the functional evidence for the operation of SOS1 exchangers in the mature root zone.

## Results

### Radioisotope imaging visualized phloem unloading of solutes in roots

To visualize the movement of solutes via the phloem using Real-time Radioisotope Imaging System (RRIS), we applied nine types of radioisotopes (^14^CO_2_ (gas), ^22^Na^+^, ^28^Mg^2+^, ^32^P-phosphate, ^35^S-sulfate, ^42^K^+^, ^45^Ca^2+^, ^59^Fe^3+^, and ^137^Cs^+^) to a leaf of Arabidopsis seedlings (Figure 1A). The distribution pattern of the radioisotope signals in the root fell into four different groups (Video 1): (1) signals were stronger in the root apex than in the mature part of the root for ^14^C, ^28^Mg, ^32^P, and ^35^S (Pattern 1; Figure 1B); (2) signals were homogenous along the root for ^42^K and ^137^Cs (Pattern 2; Figure 1C); (3) no root signal was detected for ^45^Ca and ^59^Fe (Pattern 3; Figure 1D); and (4) temporal root signals were detected for ^22^Na (Pattern 4; Figure 1E).

**Figure 1.**
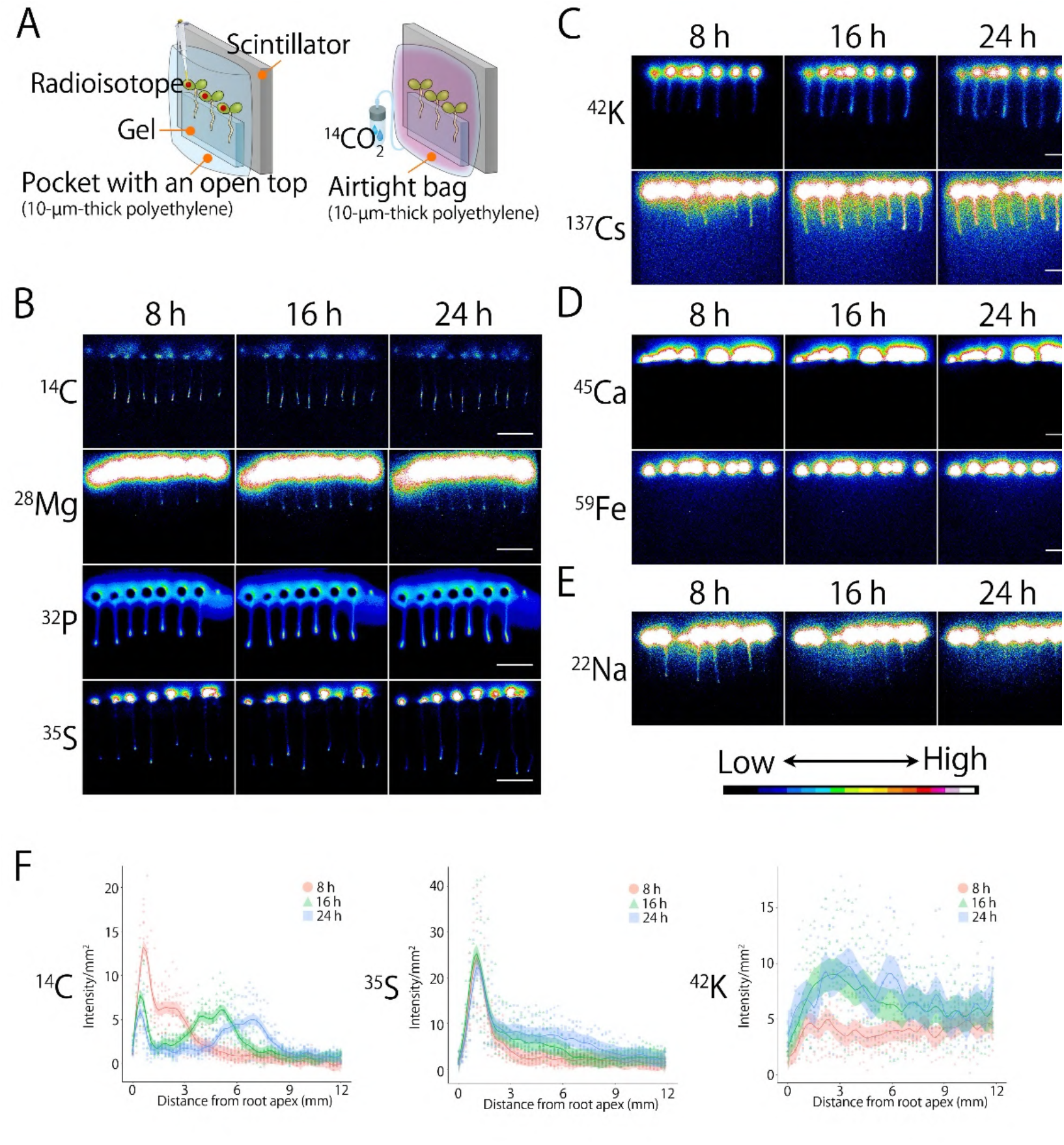
Radioisotope imaging visualizes phloem unloading of solutes in roots. Solute distribution via root phloem falls into four patterns. (**A**) Schematic diagram illustrating the application of radioisotopes to plants in radioisotope imaging using Real-time Radioisotope Imaging System (RRIS). Seven-day-old Arabidopsis seedlings were arranged on a solid medium, and radioisotopes were applied to leaves in solutions or gas (^14^CO_2_). Radiations emitted from the radioisotopes were converted into visible light, which was captured by a CCD camera to obtain images. (**B**-**E**) RRIS images of ^14^C, ^28^Mg, ^32^P, ^35^S (**B**); ^42^K, ^137^Cs (**C**); ^45^Ca, ^59^Fe (**D**); and ^22^Na (**E**) at 8, 16, and 24 h after the radioisotope application. Scale bars = 20 mm. (**F**) Radioactivity distribution profiles analyzed from the RRIS images at 8, 16, and 24 h after the application of ^14^C, ^35^S, and ^42^K. The scatter plots were made with ggplot2, R with the smooth lines by geom_smooth() command (span =0.1).

In the radioisotopes categorized as Pattern 1 (root tip accumulation), ^28^Mg, ^32^P, and ^35^S signals had a single peak at the root apex, while the ^14^C signal had two peaks around the root apex, including the vicinity of the apex (2 – 3 mm at 8 h, 4 – 5 mm at 16 h, and 5 – 7 mm at 24 h), which was likely to be the apex when ^14^C was supplied (Figure 1F). The accumulation site of ^14^C, ^32^P, and ^35^S around the root apex was further investigated using Micro-RRIS (Figure 2A). For all the three radioisotopes, the signal peak was found around 250 – 400 μm from the apex (Figure 2B). Pattern 2 radioisotopes, ^42^K and ^137^Cs, were distributed uniformly along the root throughout the observation, and the signal intensity gradually increased over time (Figure 1F). Pattern 3 radioisotopes, ^45^Ca and ^59^Fe, were not detected in the root by RRIS (Figure 1D). For Pattern 4 radioisotope ^22^Na, signal intensity in roots increased within 13 h and decreased eventually (Video 1, Figure 1E). To examine possible radioisotope extrusion from roots, we analyzed the radioactivity in the growth media after the RRIS experiment. For ^45^Ca and ^22^Na, the radioactivity of the growth media slightly increased (Supplemental Figure S1), indicating that ^45^Ca and ^22^Na were both excluded from roots to the growth media, though temporal ^45^Ca existence in the root was too scarce to detect.

**Figure 2.**
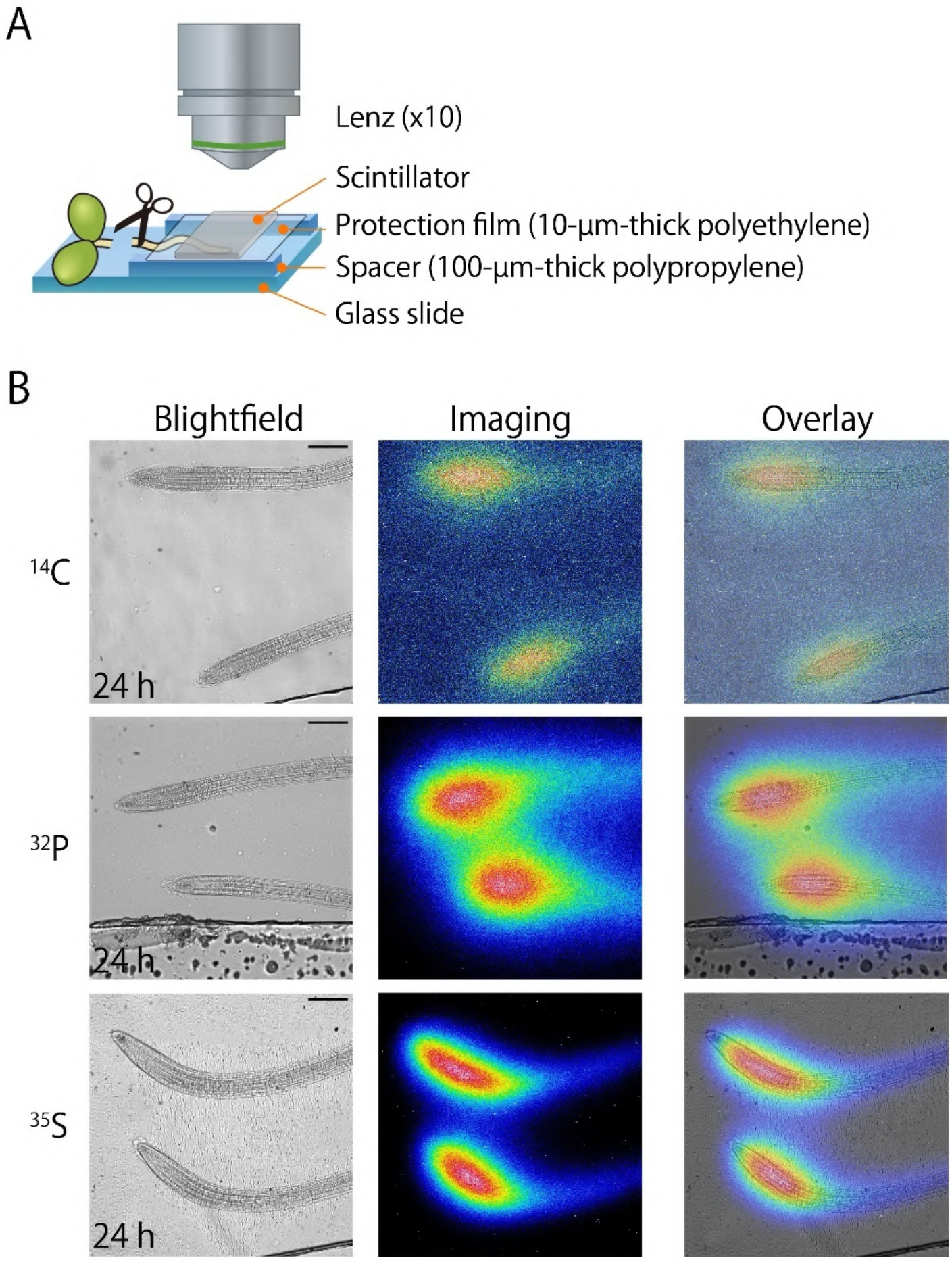
Solutes accumulate around the protophloem unloading zone in the root apex. (**A**) Schematic diagram illustrating the imaging with Micro-RRIS. Spacers were installed to prevent the roots from being deformed by the scintillator. To stop the root growth, the shoot was cut before the imaging. (**B**) Micro-RRIS images at 24 h after the foliar application of ^14^C, ^32^P, and ^35^S to seven-day-old Arabidopsis wild-type seedlings. Scale bars = 200 μm.

### Mutations in *SOS* genes caused root sodium accumulation

Root exclusion of Na^+^ uptake is mediated by the plasma membrane Na^+^/H^+^ antiporter SOS1 (Wu et al., 1996; Shi et al., 2000, 2002). However, the involvement of SOS1 in root exclusion of shoot-derived Na^+^ has not been documented. To test if SOS1 is also responsible for root exclusion of shoot-derived Na^+^, we conducted live imaging experiments using *SOS1* knockout mutants (*sos1*). After ^22^Na foliar application, we detected increasing ^22^Na signals in *sos1* roots (Video 2, Figure 3A). In contrast to the ^22^Na signals in wild-type roots, ^22^Na signals in *sos1* roots remained during 24 h of the imaging (Video 2, Figure 3A). We also conducted the imaging experiments with mutants lacking the protein kinase SOS2 (Zhu et al., 1998; Liu et al., 2000) or the Ca^2+^ sensor SOS3 (Liu and Zhu, 1997, 1998), both of which are components of the SOS pathway and essential for activation of SOS1 (Qiu et al., 2002; Quintero et al., 2002). In both *sos2* and *sos3,* ^22^Na remained in roots for 24 h (Supplemental Figure S2).

**Figure 3.**
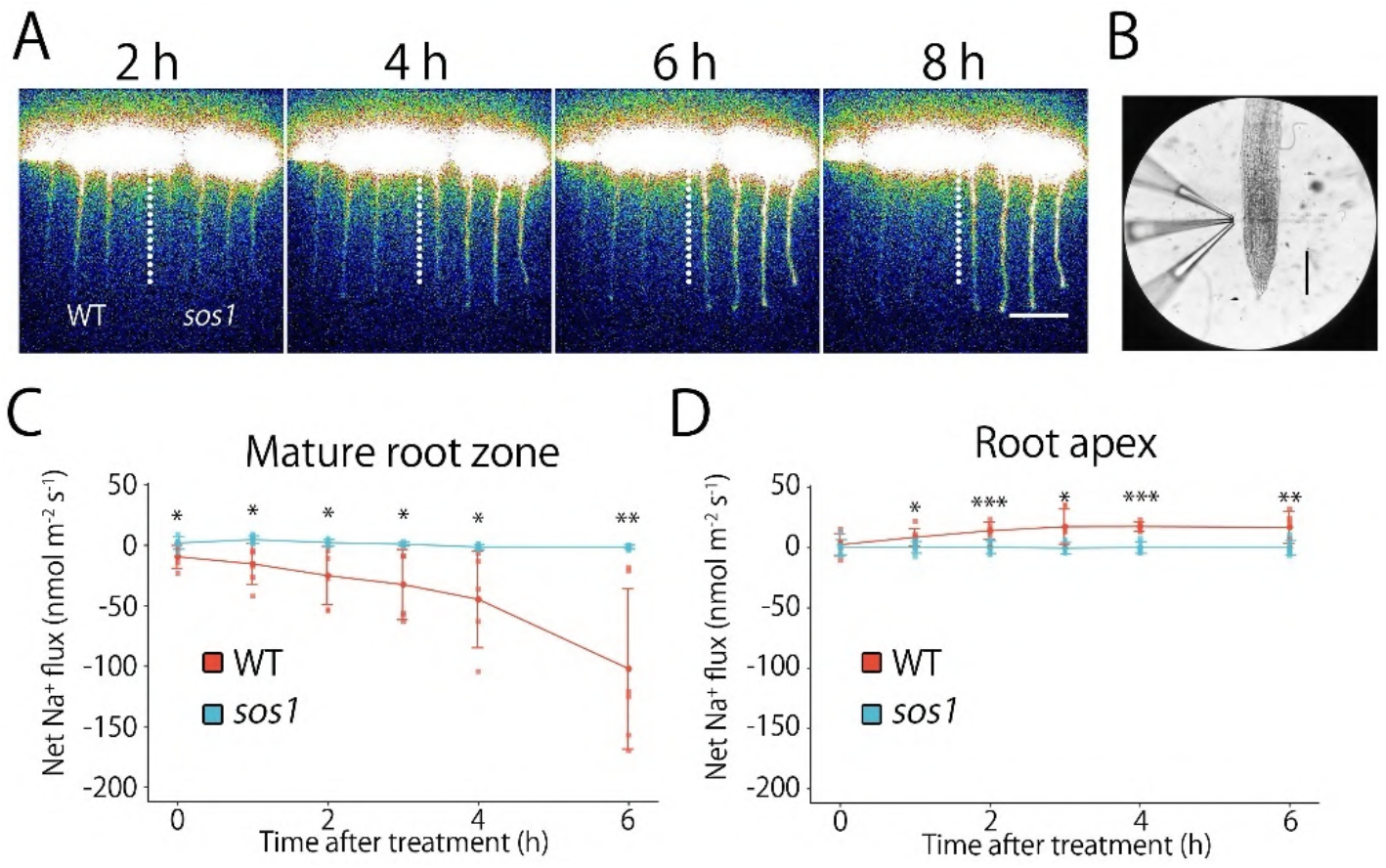
Sodium-ion exclusion is active at the mature root zone. (**A**) RRIS images at 2, 4, 6, and 8 h after foliar application of ^22^Na to seven-day-old Arabidopsis seedlings of the wild type (WT, left) and *SOS1* knockout mutant (*sos1*, right). Scale bars = 20 mm. (**B**) Positioning of microelectrodes during the ion flux measurement by Microelectrode Ion Flux Estimation (MIFE) technique. The tips of three ion-selective microelectrodes for Na^+^, K^+^, and H^+^ were positioned close to each other at 40 μm apart from the root surface. Scale bar = 200 mm. (**C, D**) Net Na^+^ flux from the surface of the mature root zone (**C**, 5 mm from the root cap) and root apex (**D**, 400 – 600 μm from the root cap) after the foliar application of 5 mM NaCl solutions to seven-day-old Arabidopsis seedlings of WT (red) and *sos1* (blue). The net flux value of each replicate (*n* = 5 – 7) is represented as a dot with means and standard deviations. Asterisks indicate a significant difference in net flux between WT and *sos1* (Student’s *t*-test, *\*P* < 0.05, *\*\*P* < 0.01, \*\*\**P* < 0.001).

### Sodium exclusion was active at the mature root zone

To determine the root zones of Na^+^ exclusion, we analyzed patterns of net ion fluxes at the surface of root tissue using the non-invasive Microelectrode Ion Flux Estimation (MIFE) technique (Figure 3B; Shabala et al., 1997; Newman, 2001). Following foliar application of NaCl solution, the magnitude of Na^+^ efflux increased within 6 h at the mature zone of wild-type roots, while such net efflux was not observed in *sos1* roots (Figure 3C). Continuous measurement of net Na^+^ flux in wild-type roots demonstrated that the magnitude of Na^+^ efflux increased until 12 h (Supplemental Figure S3). We did not detect such Na^+^ efflux when we applied sorbitol solution causing the same level of osmotic pressure as the NaCl solution (Supplemental Figure S4), indicating that the increasing Na^+^ efflux was NaCl-specific but not triggered by osmotic stress. In the presence of amiloride, which is an inhibitor of Na^+^/H^+^ antiporters, including SOS1 (Wu et al. 2019), wild-type roots did not show Na^+^ efflux (Supplemental Figure S4). At the root apex, net Na^+^ flux of wild-type roots showed net Na^+^ influx (Figure 3D), possibly representing reabsorption of the excluded Na^+^, considering Na^+^ was not supplied in the medium.

Simultaneously analyzed net K^+^ flux and net H^+^ flux showed no significant difference between plants with/without Na^+^ efflux activity (Supplemental Figure S5). We further analyzed the electrical potential difference across the plasma membrane at the mature root zone of wild-type seedlings. The foliar application of NaCl solution caused the hyperpolarization of root epidermal cells after 6 h (Supplemental Figure S6), which was after the magnitude of Na^+^ efflux started to increase.

## Discussion

### Sodium exclusion from roots

The live imaging of radiotracers and net ion flux analysis demonstrated root exclusion of Na^+^ following the application of NaCl solution to Arabidopsis leaves (Video 1, Figures 1E, 3C). Using *SOS* mutants and SOS1 inhibitor, the SOS pathway was shown to be essential for this root Na^+^ exclusion (Video 2, Figures 3A, 3C, Supplemental Figure S2), which provides the first evidence of SOS pathway involvement in the root exclusion of shoot-derived Na^+^. The ^22^Na accumulation in *SOS* mutant roots also suggests that the SOS pathway is not essential for Na^+^ exclusion from the shoot.

In studies employing the electrode technique for ion flux analyses, net Na^+^ flux data were often regarded as unreliable because of a technical problem that commercially available Na^+^ ionophore cocktails had poor Na^+^ selectivity over K^+^ and Ca^2+^ (Knowles and Shabala, 2004; Chen et al., 2005; Shabala et al., 2005). However, a newly developed calixarene-based Na^+^ ionophore cocktail, which we used in the present study, has shown better Na^+^ selectivity over K^+^ and Ca^2+^ (Carden et al., 2001; Jayakannan et al., 2011). We also confirmed electrode Na^+^ selectivity over other ions before each of our ion flux measurements. Simultaneously analyzed net Na^+^ flux and net K^+^ flux showed a clear difference in patterns (Figures 3C, 3D, Supplemental Figure S5), providing another example that demonstrates the reliability of the newly developed Na^+^ ionophore cocktail.

The expression pattern of *SOS1* was previously investigated with transgenic Arabidopsis plants harboring the *Escherichia coli* β-glucuronidase (GUS) gene under the control of *SOS1* promoter, which showed strong GUS expression at the root tip epidermis (Shi et al., 2002). In the mature root zone, weaker GUS expression was observed in the xylem parenchyma cells (Shi et al., 2002), which are located inside the Casparian strips. Based on this expression pattern, root Na^+^ exclusion by SOS1 has been regarded as active at the root apex. However, cell-type-specific microarray expression profile in Arabidopsis roots showed *SOS1* expression in nearly all root cell types, including outer layers of the mature root zone (Brady et al., 2007). Consistently, a previous net ion flux analysis indicated that *SOS1* knockout mutation affects the ion flux kinetics in both the root apex and mature root zone (Shabala et al., 2005). The present study suggested that SOS1 mediates the exclusion of shoot-derived Na^+^ in the mature root zone (Figures 3C, 3D). This might prevent Na^+^ from reaching meristematic cells, which lack vacuoles to sequestrate Na^+^. Also, considering plant Na tolerance can be correlated to the capacity of root Na^+^ exclusion (Lessani and Marschner, 1978), it is reasonable that Na^+^ is excluded from the mature root zones, which account for a large fraction of the root. To confirm the SOS1 function in the mature root zone, SOS1 protein localization needs to be examined.

Activation of SOS1 through the SOS pathway was essential for root Na^+^ exclusion we observed, based on the inhibition of root Na^+^ exclusion in *sos2* and *sos3* (Supplemental Figure S2). SOS1 is activated by phosphorylation of an autoinhibitory domain by the protein kinase SOS2 (Quintero et al., 2011; Jarvis et al., 2014). SOS2 is activated through interaction with Ca^2+^ sensors such as SOS3 (Halfter et al., 2000) and SOS3-like calcium-binding protein 8 (SCaBP8, Quan et al., 2007; Kim et al., 2007). As a trigger of the SOS pathway, cytosolic free Ca^2+^ concentration increases in response to salt stress (Knight et al., 1997). Its underlying mechanism is explained by Na^+^ binding to glycosyl inositol phosphorylceramide (GIPC) sphingolipids at the cell surface to gate Ca^2+^ influx channels (Jiang et al., 2019). Interestingly, in the current study, we observed SOS pathway-involved root Na^+^ exclusion in the absence of severe salt stress: Na^+^ concentrations of the growth media (2 mM) and the solutions applied to leaves (0.5 mM) were both in a low range. This indicates that either the SOS pathway can be triggered by a subtle increase in cytosolic Na^+^ concentration through an unidentified mechanism of Ca^2+^ signaling, or SOS1 is activated through the SOS pathway constantly under normal conditions and there is a basal level of active SOS1 to mediate Na^+^ exclusion.

### Variation of phloem unloading manner

Accumulation of Pattern 1 radioisotopes (^14^C, ^28^Mg, ^32^P, and ^35^S) in the root apex (Figure 1B) suggested that most of the compounds containing those elements are not unloaded from the phloem until they reach the terminal region of the root (Figure 4). The accumulation peak of ^14^C, ^32^P, and ^35^ S in the root apex was around 250 – 400 μm from the root cap (Figure 2B), which corresponded to the location of the terminal protophloem sieve elements (Ross-Elliott et al., 2017). Among those elements, only ^14^C had another peak in the upper part of the root (Figure 1F), suggesting that ^14^C unloading at the root apex is more active compared to the other Pattern 1 elements (Figure 4). The difference in the unloading activity is understandable, considering there would be a higher demand for carbon in the root apex for cell division and extension.

**Figure 4.**
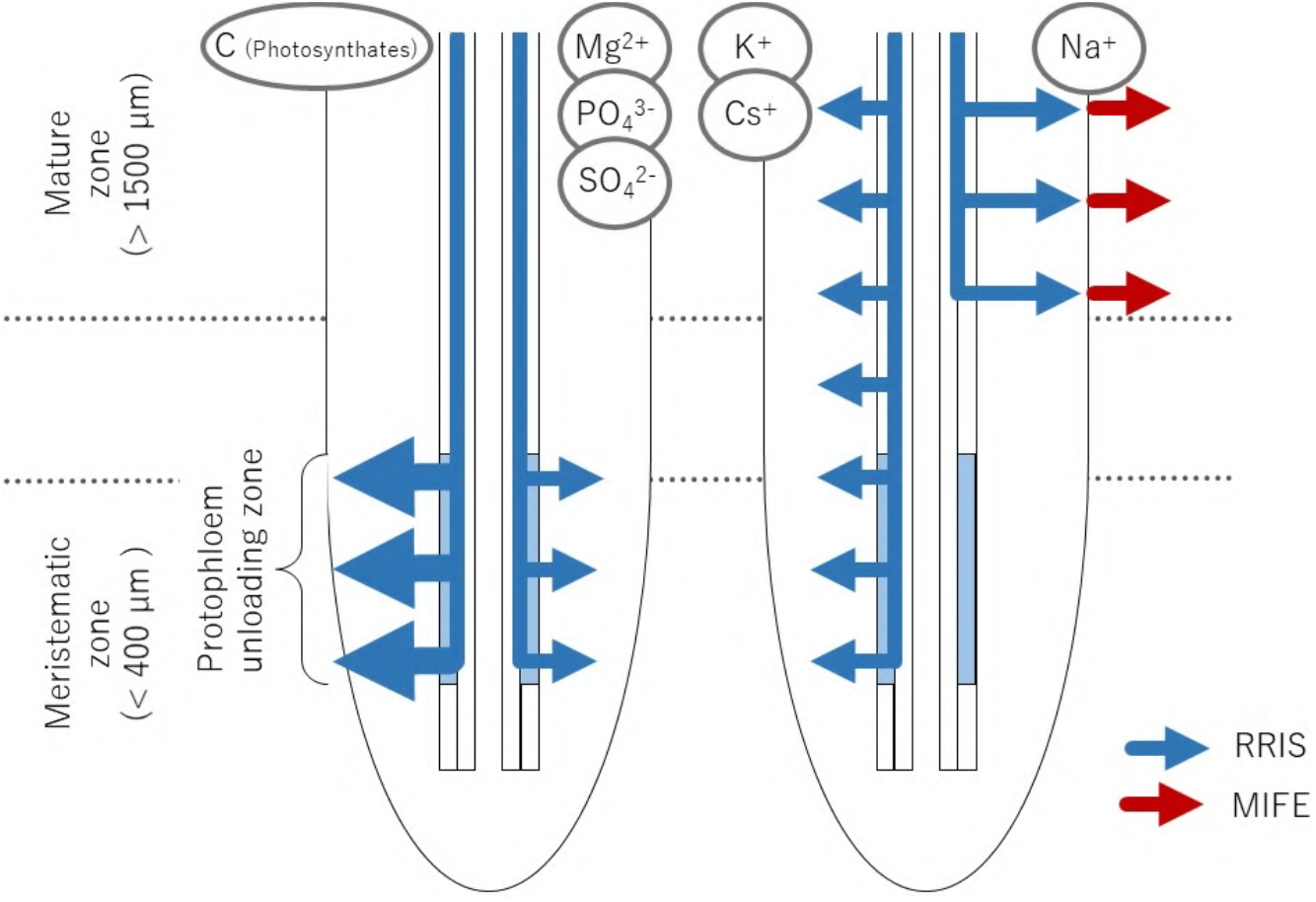
Schematic model of the solute movement via the phloem in Arabidopsis roots. Phloem unloading of C (Photosynthates), Mg, P, and S is inactive in the mature root zone and they reach the root apex once in the phloem flow. Among these elements, unloading in the meristematic zone is more active for C. For Mg, P, and S, elements reaching the apex move downwards accordingly with the root elongation. In contrast, K and Cs are actively unloaded not only in the meristematic zone but also in the mature zone. Na is unloaded from the phloem and excluded into the rhizosphere in the mature zone by the operation of SOS1 Na^+^/H^+^ exchangers. The red and blue arrows indicate the solute movement confirmed by the radioisotope imaging (RRIS) and the ion flux analysis (MIFE), respectively.

In contrast, Pattern 2 radioisotopes (^42^K and ^137^Cs) were unloaded in mature root zones, and there was no prominent peak at the root apex (Figures 1C, 1F), suggesting that ion channels or transporters are functioning in the mature root zone to actively unload K^+^ and Cs^+^ from the phloem (Figure 4). Potassium is necessary for all types of cells because K^+^ acts as a coenzyme that promotes various physiological processes in the plant cell (Taiz and Zeiger, 2002), which might be a reason why K^+^ was actively unloaded from the phloem.

In radiotracer analyses, conventional autoradiography has visualized elemental distribution in fixed samples at a single time point after the addition of radiotracers. With those methods, it is difficult to keep track of changes in elemental distribution over time. To overcome this problem, we have developed a live imaging technique that facilitates the kinetic analysis of various nuclides (^14^C, ^22^Na, ^28^Mg, ^32^P, ^35^S, ^36^Cl, ^42^K, ^45^Ca, ^54^Mn, ^59^Fe, ^65^Zn, ^86^Rb, ^109^Cd, and ^137^Cs). In the present study, we utilized the live imaging technique and presented novel findings such as the changes in distribution patterns of ^14^C-photosynthates along the root (Figure 1E) and the transient accumulation of ^22^Na in roots (Figure 1F), which would be difficult to find with conventional autoradiography. Furthermore, we present an example where limitations in traceability and spatial resolution of autoradiography can be overcome by the complementary use of non-invasive ion flux analysis, which in our case identified the location of Na^+^ exclusion from the root.

## Materials and methods

### Plant materials and growth conditions

Seeds of *Arabidopsis thaliana* ecotype Columbia-0 (Inplanta Innovations Inc., Yokohama, Japan) and T-DNA insertion mutant lines of *AtSOS1, AtSOS2,* and *AtSOS3* (CS3862, CS3863, and CS3864, respectively; all from Arabidopsis Biological Resource Center, Columbus, OH, USA) were surface sterilized with 5% sodium hypochlorite for 5 min and rinsed thoroughly with distilled water. For radiotracer experiments, plants were grown on MGRL medium (Fujiwara et al., 1992) solidified with 0.4% gellan gum (FUJIFILM Wako Pure Chemical, Osaka, Japan) at 22°C under a photoperiod of 16 h light (100 μmol m^-2^ s^-1^) and 8 h darkness for seven days. For ion flux analyses, plants were grown under the same conditions for seven days except for modifications in the growth medium: to exclude Na^+^ from the medium, 1.5 mM NaH2_P_O_4_ and 0.25 mM Na_2_HPO_4_ in the original MGRL medium were replaced with 1.75 mM (NH_4_)_2_PO_4_, and 67 μM EDTA-2Na and 8.6 μM FeSO_4_ in the original medium were replaced with 65 μM Fe(III)-EDTA (Fe^3+^ 8.6 μM). With the modifications, KOH (ca. 250 μM) was added to maintain the pH at 5.8 because the phosphate functioning as a buffer in the original medium was removed.

### Radioisotope imaging of phloem unloading in roots

To visualize phloem unloading, we used Real-time Radioisotope Imaging System (RRIS; Nakanishi et al., 2009). Seven-day-old Arabidopsis seedlings were transferred onto a new solid growth medium with the shoot hanging on the edge of the medium. To bring the plant roots into contact with a fiber optic plate with a scintillator (FOS), plants on the medium were placed into a 10-μm-thick polyethylene pocket with an open top, and the pocket was placed on the FOS in the direction that the roots came between the medium and FOS (Figure 1A). For radioisotopes except ^14^C, 1 μl of radioisotope solution was dropped onto a cotyledon. For the imaging of ^14^C, the opening top of the polyethylene pocket was sealed to make an airtight bag. ^14^CO_2_ was produced by mixing [^14^C]-sodium bicarbonate (5 MBq) and lactic acid in a 1.5 ml vial with a septum stopper equipped with a syringe needle (Sugita et al. 2013), and the produced ^14^CO_2_ was pumped into the airtight bag containing plants by connecting the vial and bag with a needle and tube. Imaging started immediately after radioisotope applications in a dark box. Images were taken hourly by CCD camera (C3077-70, Hamamatsu Photonics Co., Shizuoka, Japan) for 15 min in the darkness, and plants were illuminated during the other 45 min, as previously described by Hirose et al. (2013). The cycle of imaging and illumination was repeated until 24 h after the application of radioisotopes. For the imaging of ^14^C, the airtight bag was opened at 5 h to prevent CO_2_ deficiency since the CO_2_ concentration around 400 ppm at the start of the imaging decreased in time to leave little ^14^CO_2_ signals in the air after 5 h. Concentrations of radioisotopes and carrier ions in radioisotope solutions were respectively as follows: ^28^Mg^2+^, 5 kBq μl^-1^ and 1 mM; ^32^P-phosphate, 5 kBq μl^-1^ and 0.5 mM; ^35^S-sulfate, 50 kBq μl^-1^ and 1 mM; ^42^K^+^, 3 kBq μl^-1^ and 15 mM; ^137^Cs^+^, 3.5 kBq μl^-1^ and 1 mM; ^45^Ca^2+^, 50 kBq μl^-1^ and 1 mM; ^59^Fe^3+^, 10 kBq μl^-1^ and 1 mM; ^22^Na^+^, 3.5 kBq μl^-1^ and 0.5 mM. ^28^Mg^2+^ was produced by ^27^Al(α, 3p)^28^Mg reaction and was separated from the Al target (Iwata et al., 1992). ^42^K^+^ was prepared from a ^42^Ar-^42^K generator by milking (Aramaki et al., 2015). [^14^C]-sodium bicarbonate, ^32^P-phosphate, ^35^S-sulfate, ^45^Ca^2+^ and ^59^Fe^3+^ were purchased from PerkinElmer (Waltham, MA, USA). ^137^Cs^+^ and ^22^Na^+^ were purchased from Eckert & Ziegler (Valencia, CA, USA). For the acquired images, the radioisotope distribution profile along the main root was analyzed using Image J software.

For the imaging of the root tip area using Micro-RRIS, plants were taken from the polyethylene pocket after 24 h of live imaging. The plants were placed on a glass slide, and the FOS with a protection film was placed on the root, with spacers pre-installed in between the glass slide and FOS to prevent the root from being crushed (Figure 2A). Images were taken in the darkness by CCD camera (iXon 888, Andor Technology Ltd., Belfast, UK)) with an exposure time of 5 min for ^32^P and 60 min for ^14^C and ^35^S.

### Quantification of radioisotope distribution

To quantify the distribution of ^45^Ca and ^22^Na after the foliar application, seven-day-old Arabidopsis seedlings were set on a solid growth medium with a size of 30 x 50 mm in the same manner as the real-time imaging, and 1 μl of radioisotope solution was dropped onto a cotyledon, which we refer to as the labeled cotyledon. Plants were incubated in a small chamber at 22°C under the light (100 μmol m^-2^ s^-1^). After 24 h, plants were divided into four parts (the labeled cotyledon, the other cotyledon, hypocotyl, and root), and the radioactivity of ^45^Ca and ^22^Na in each part and the medium was measured with a liquid scintillation counter (LSC-6100, Hitachi, LTD. Tokyo, Japan) and a gamma counter (ARC-300, Hitachi, LTD. Tokyo, Japan), respectively.

### Determination of net ion flux in roots

Net ion flux at the surface of the root was measured non-invasively with ion-selective microelectrodes (MIFE technique; Shabala et al., 1997; Newman, 2001). Ion-selective microelectrodes and the reference electrode were fabricated as previously described (Shabala et al., 1997). For the preparation of Na^+^-, K^+^-, and H^+^-selective microelectrodes, microelectrodes were backfilled with 500 mM NaCl, 200 mM KCl, and 15 mM NaCl plus 40 mM KH_2_PO_4_, respectively. Then Na^+^-selective microelectrodes were front-filled with an improved calixarene-based Na^+^ ionophore cocktail (Jayakannan et al., 2011), and K^+^- and H^+^-selective microelectrodes were front-filled with commercially available ionophore cocktails (Product numbers 60031 and 95297, respectively; Sigma-Aldrich, St. Louis, MO, USA). Three microelectrodes were mounted on a 3D-micromanipulator (MMT-5, Narishige, Tokyo, Japan), and calibrated using sets of three solutions with different ion concentrations (10, 20, 50 μM NaCl for Na^+^; 10, 20, 50 μM KCl for K^+^; pH 5.1, 6.4, 7.8 for H^+^). Only electrodes with a Nernstian slope of more than 50 mV/decade and a correlation of more than 0.999 were used. The tips of the microelectrodes were positioned close together, 40 μm above the root surface. A computer-controlled stepper motor moved the microelectrodes every 6 sec (or every 30 sec for continuous measurements) between two positions, 40 μm and 150 μm from the surface of the sample root. The potential difference between the two positions was recorded, and net ion flux was calculated as previously described (Newman, 2001). For continuous measurements, average net flux values for every 5 min were calculated.

To localize the site of Na^+^ exclusion, Na^+^ flux was determined at the root apex (400 – 600 μm from the root cap) and the mature root zone (5 mm from the root cap) of Arabidopsis wild type and *sos1* seedlings. Seven-day-old Arabidopsis seedlings were immobilized on a microscope slide using strips of a paraffin film, and the root was pretreated with distilled water or 100 μM amiloride solution (Na^+^ 0 μM for both conditions), with the shoot kept above the solution. To stabilize biological activities after the possible disturbance from the immobilization, the immobilized root was kept in the pretreatment solutions for 1 h. Before the treatment, net ion (Na^+^, K^+^, and H^+^) flux from the root apex and the mature root zone was measured for 3 – 5 min to ensure steady initial values. A 5 μL droplet of 5 mM NaCl solution or 8.5 mM sorbitol solution was applied onto a cotyledon, and net ion flux from the two zones was measured hourly until 6 h after the application. Five to seven replicates were tested for each condition and line.

### Determination of membrane potential

The electrical potential difference across the plasma membrane was determined by measuring electrical potentials inside and outside the plasma membrane using a microelectrode as previously described (Shabala and Newman, 1999). For the microelectrode preparation, a borosilicate glass capillary with an internal filament (GC150F-10, Harvard Apparatus, Cambridge, MA, USA) was pulled and filled with 500 mM KCl solution. A reference electrode was prepared in the same manner as the net ion flux analysis. Seven-day-old Arabidopsis wild-type seedlings were immobilized on plastic blocks using strips of a paraffin film, and the roots were kept in 10 mL of distilled water for 1 h. A 5 μL droplet of 5 mM NaCl solution was applied onto a cotyledon, and the electrical potential in an epidermal cell was measured by impaling the cell with the microelectrode. Membrane potential was measured on three occasions: before the NaCl application, and 1 h and 6 h after the NaCl application. Average membrane potential values were calculated from two to six impalements made for each seedling, and four to six seedlings were tested for each condition.

### Statistical analysis

The Student’s *t*-test was carried out using R software version 4.1.1 to compare differences in net Na^+^ flux between the wild type and *SOS1* knockout mutant. A factorial analysis of variance (ANOVA) was used to test differences in membrane potential of root epidermal cells at different time points. When a difference was found, the Tukey-Kramer test was carried out using R software version 4.1.1 to compare the values of different conditions and to test in which comparison the difference was significant. Significance was set at the 5% level.

## Acknowledgements

This work was supported by funding from the Japan Society for the Promotion of Science (JSPS) KAKENHI Grant No. 20K06317 to RS, 19J22719 to TO, 20H00437 to TMN, and 19KK0148 to KT.

## Author contributions

RS, Conceptualization, Acquisition of data, Formal analysis, Funding acquisition, Methodology, Resources, Visualization, Writing – original draft, Writing – review and editing; TO, Acquisition of data, Formal analysis, Funding acquisition, Methodology, Visualization, Writing – original draft, Writing – review and editing; NIK, Conceptualization, Methodology, Resources, Supervision, Validation, Writing – review and editing; MBG, Acquisition of data, Methodology, Validation; LS, Methodology, Resources, Supervision, Validation, Writing – review and editing; TMN, Funding acquisition, Resources, Validation; SS, Methodology, Resources, Supervision, Validation, Writing – review and editing; KT, Conceptualization, Formal analysis, Funding acquisition, Methodology, Project administration, Resources, Supervision, Validation, Visualization, Writing – original draft, Writing – review and editing

---

**Video 1. Animations of ^14^C, ^28^Mg, ^32^P, ^35^S, ^42^K, ^137^Cs, ^45^Ca, ^59^Fe, and ^22^Na created by RRIS.** Animations were created from a series of images taken hourly by RRIS after the foliar application of radioisotopes to seven-day-old Arabidopsis wild-type seedlings. Scale bars = 20 mm.

**Video 2. Knockout mutation in *SOS1* causes root accumulation of shoot-derived sodium.** The animation was created from a series of images taken hourly by RRIS after foliar application of ^22^Na to seven-day-old Arabidopsis seedlings of the wild type (WT, left) and *SOS1* knockout mutant (*sos1,* right). Scale bars = 20 mm.

**Supplemental Figure S1.**
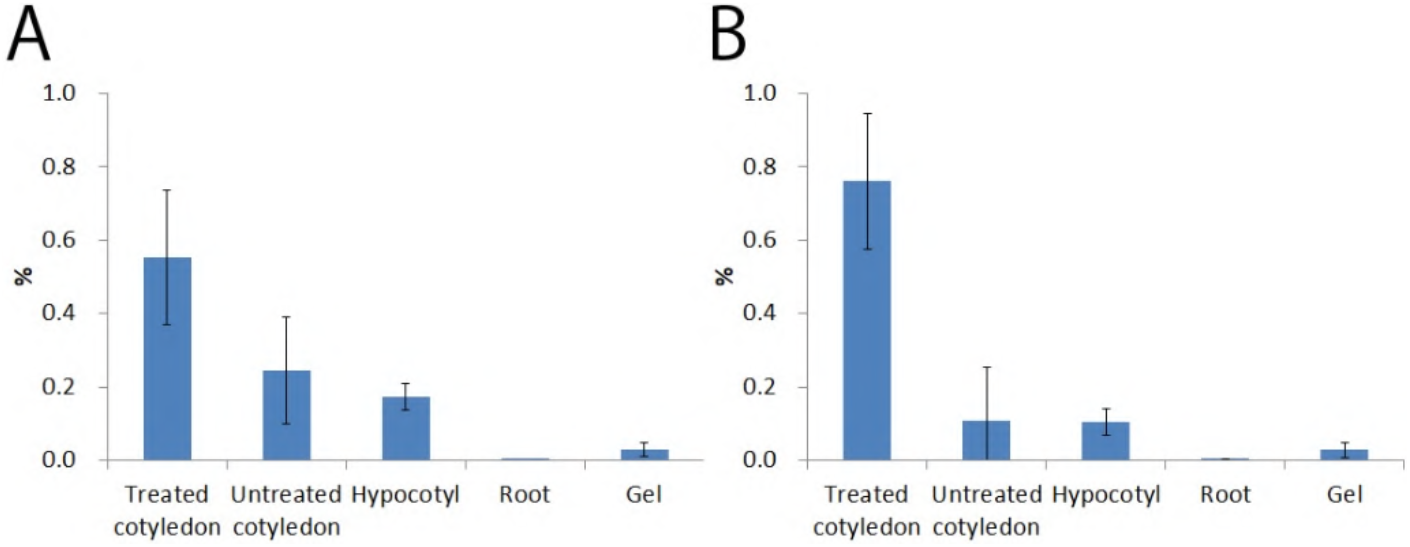
Distribution of ^45^Ca and ^22^Na after application to a cotyledon. (**A, B**) The ratio of distributed radioisotopes in each tissue or growth medium to the total radioisotopes applied to a cotyledon. Twenty-four hours after the radioisotope application to a cotyledon of seven-day-old Arabidopsis wild-type seedlings, plants were divided into four parts (the labeled cotyledon, the other cotyledon, hypocotyl, and root), and the radioactivity of ^45^Ca (**A**) and ^22^Na (**B**) in each part and the medium was determined by a liquid scintillation counter and a gamma counter, respectively.

**Supplemental Figure S2.**
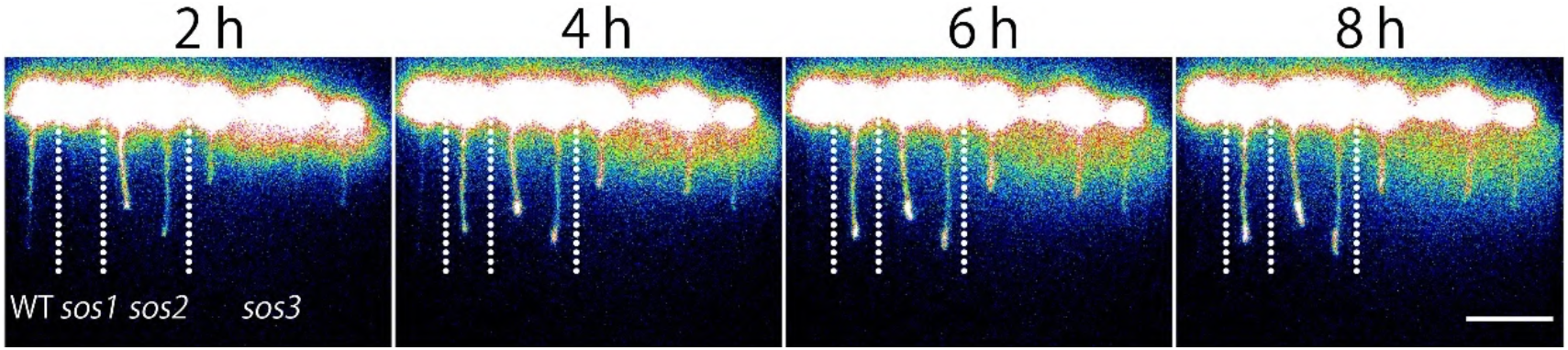
Knockout mutations in each of *SOS1, SOS2,* and *SOS3* cause root accumulation of shoot-derived sodium. RRIS images at 2, 4, 6, and 8 h after foliar application of ^22^Na to seven-day-old Arabidopsis seedlings of the wild type (WT) and single knockout mutants of *SOS1, SOS2,* and *SOS3 (sos1, sos2,* and *sos3,* respectively). Scale bars = 20 mm.

**Supplemental Figure S3.**
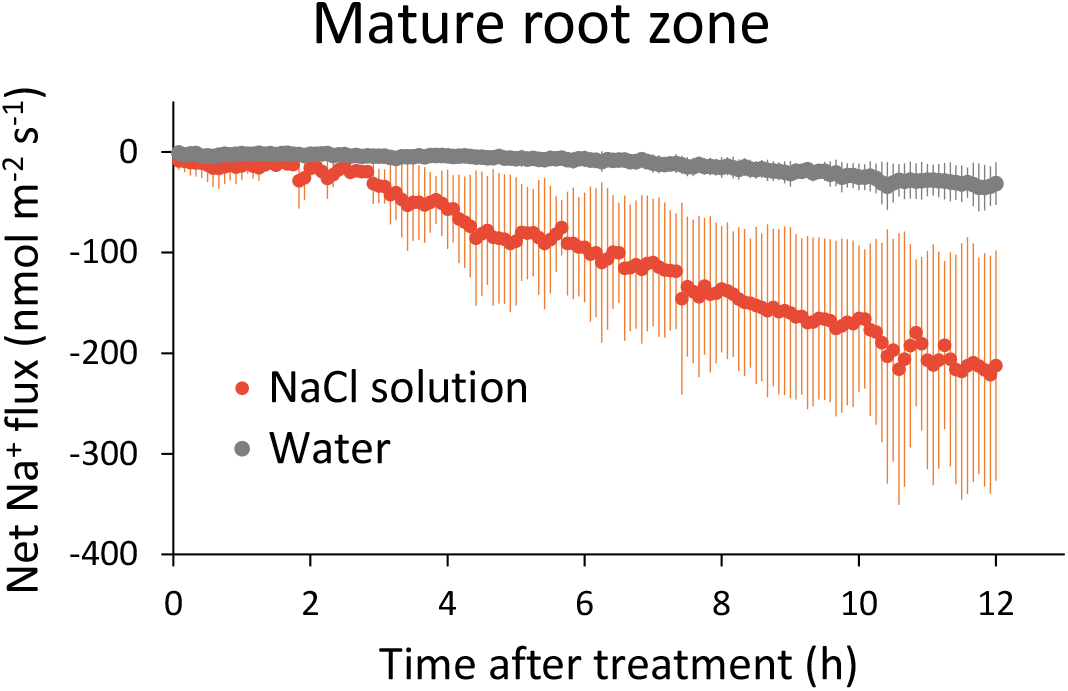
Continuous measurement of net sodium ion flux at the mature root zone. Net Na^+^ flux from the surface of the mature root zone (5 μm from the root cap) after foliar application of 5 mM NaCl solution (red) or water (grey) to seven-day-old Arabidopsis wild-type seedlings. Data represent means (NaCl solution, *n* = 3; water, *n* = 2) with standard deviation.

**Supplemental Figure S4.**
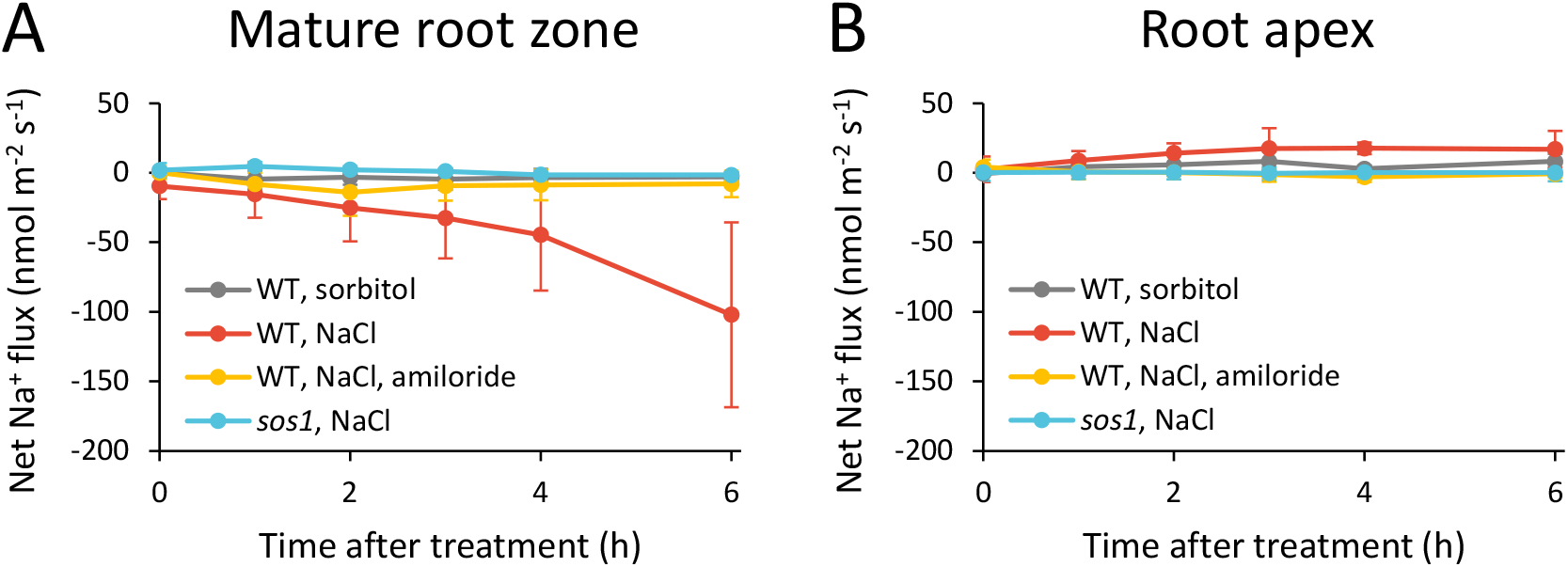
Sodium-ion exclusion at the mature root zone is not induced by osmotic stress and is inhibited by amiloride. (**A**, **B**) Net Na^+^ flux from the surface of the mature root zone (**A**, 5 mm from the root cap) and root apex (**B**, 400 – 600 μm from the root cap) of seven-day-old Arabidopsis seedlings after the foliar application of 8.5 mM sorbitol solutions to the wild type (WT, grey), 5 mM NaCl solutions to WT (red), 5 mM NaCl solutions to WT treated with 100 μM amiloride solution (yellow), and 5 mM NaCl solutions to *SOS1* knockout mutant (*sos1,* blue). Data represent means (*n* = 5 – 7) with standard deviations.

**Supplemental Figure S5.**
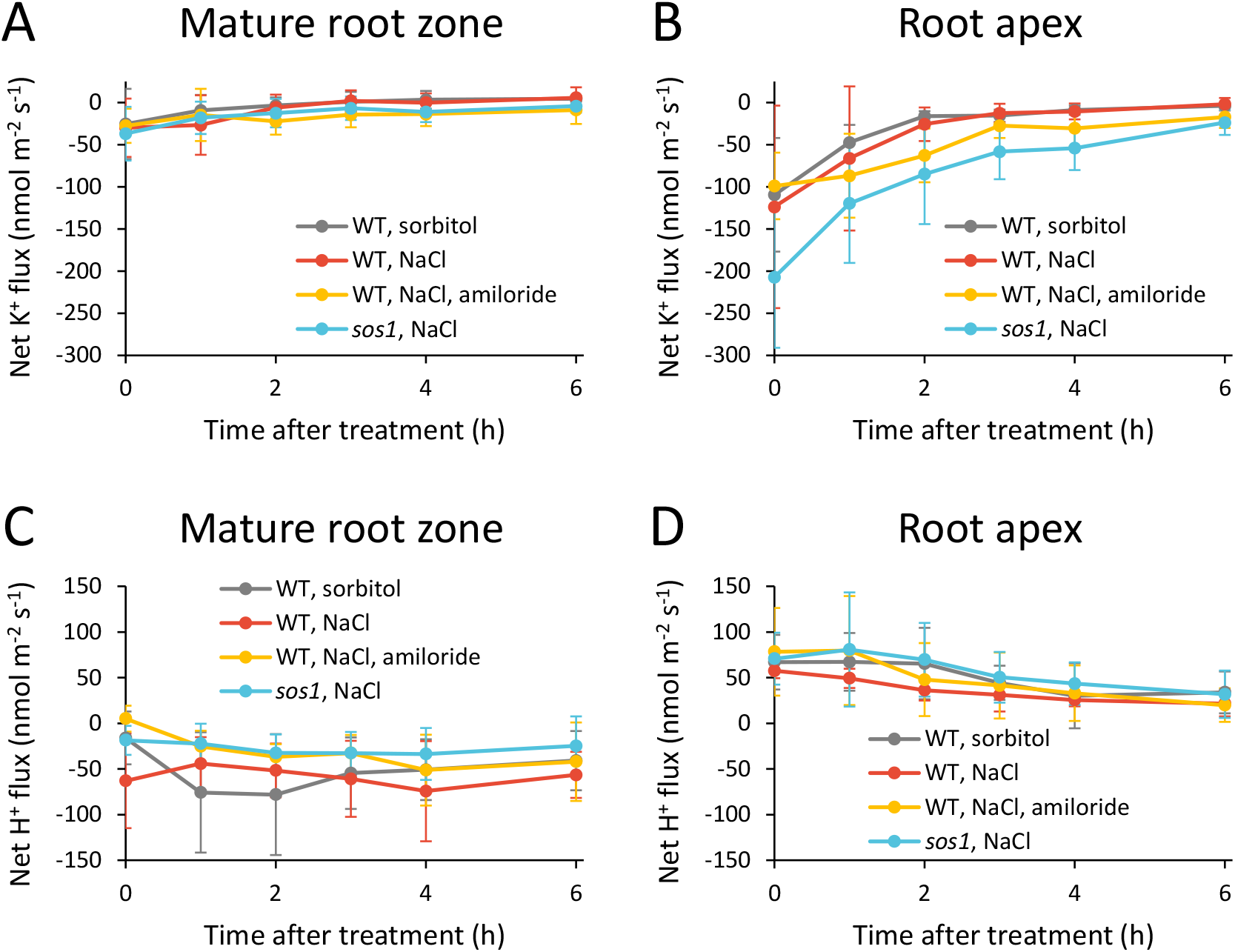
Sodium-ion exclusion is not accompanied by changes in net potassium ion flux or net proton flux. (**A-D**) Net K^+^ flux (**A**, **B**) and net H^+^ flux (**C**, **D**) from the surface of the mature root zone (**A**, **C**; 5 mm from the root cap) and root apex (**B**, **D**; 400 – 600 μm from the root cap) of seven-day-old Arabidopsis seedlings after the foliar application of 8.5 mM sorbitol solutions to the wild type (WT, grey), 5 mM NaCl solutions to WT (red), 5 mM NaCl solutions to WT treated with 100 μM amiloride solution (yellow), and 5 mM NaCl solutions to *SOS1* knockout mutant (*sos1,* blue). Data represent means (*n* = 5 – 8) with standard deviations.

**Supplemental Figure S6.**
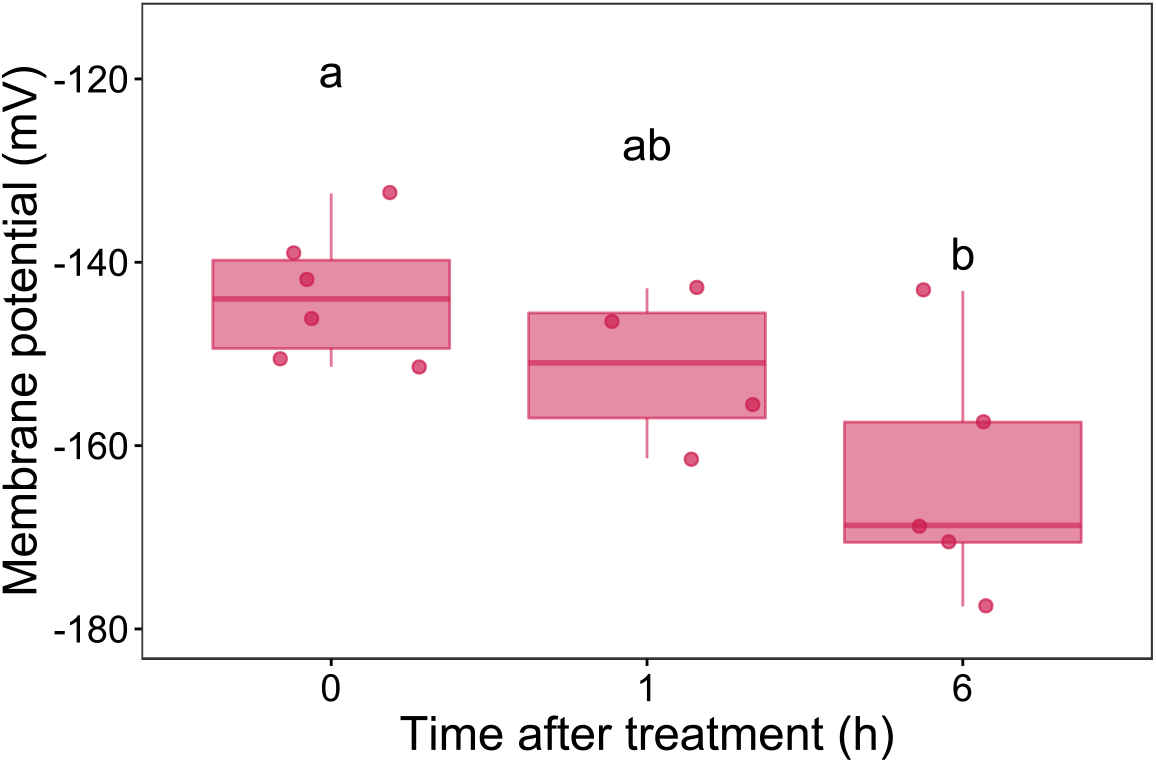
Sodium-ion exclusion is accompanied by hyperpolarization of root epidermal cells. The membrane potential of epidermal cells in mature root zone after the foliar application of 5 mM NaCl solutions to seven-day-old Arabidopsis wild-type seedlings. Membrane potential was measured before, 1 h after, and 6 h after the foliar application. The average membrane potential value calculated from repeated measurements in each replicate is represented as a dot. Box plots show median (*Q_2_*), first (*Q_1_*) and third (*Q_3_*) quartile, minimum (*Q_0_*), and maximum (*Q_4_*) (*n* = 4 – 6). Different letters indicate a significant difference (one-way ANOVA and Tukey’s HSD test, p < 0.05).

