## Supplementary figures and images for "Isotope-based visualization of element distribution in phloem provides functional evidence for the operation of SOS1 Na^+^/H^+^ exchangers in mature zones of Arabidopsis root"

### Supplemental Figure S1

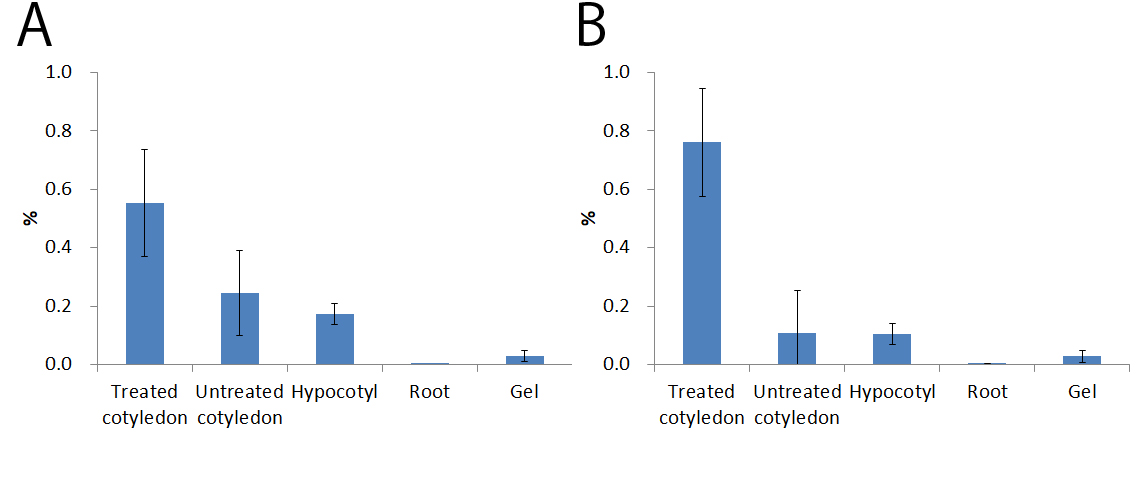

### Supplemental Figure S2

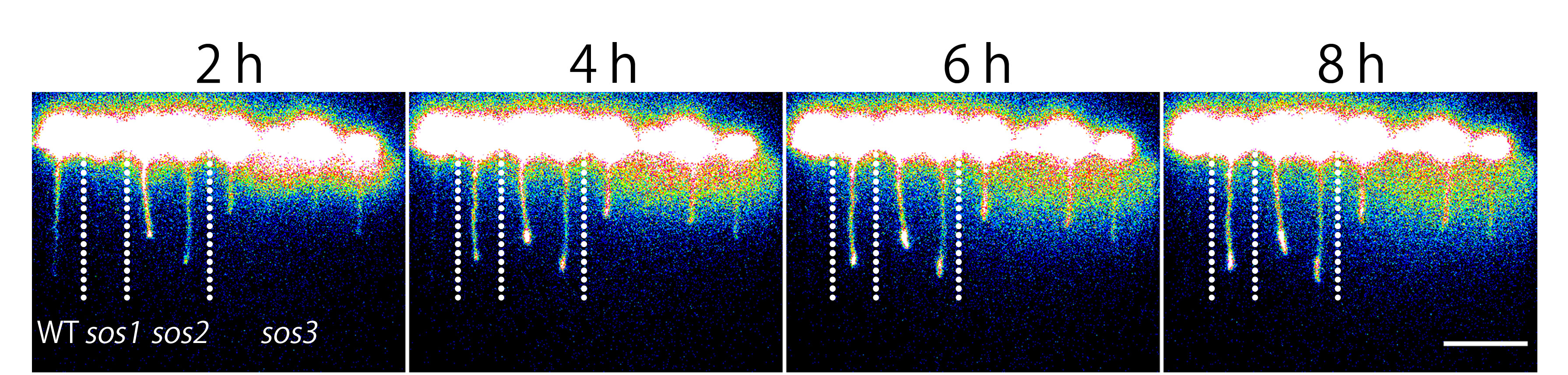

### Video 1

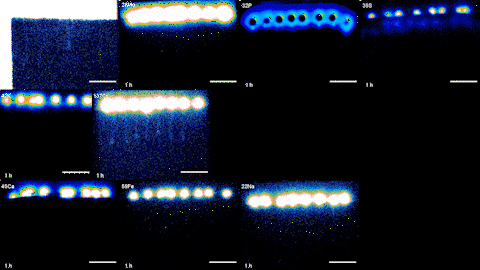

### Video 2

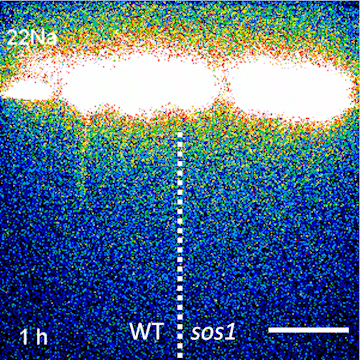
