## Supplemental Figure S3-S6 for "Isotope-based visualization of element distribution in phloem provides functional evidence for the operation of SOS1 Na^+^/H^+^ exchangers in mature zones of Arabidopsis root"

### Slide 1
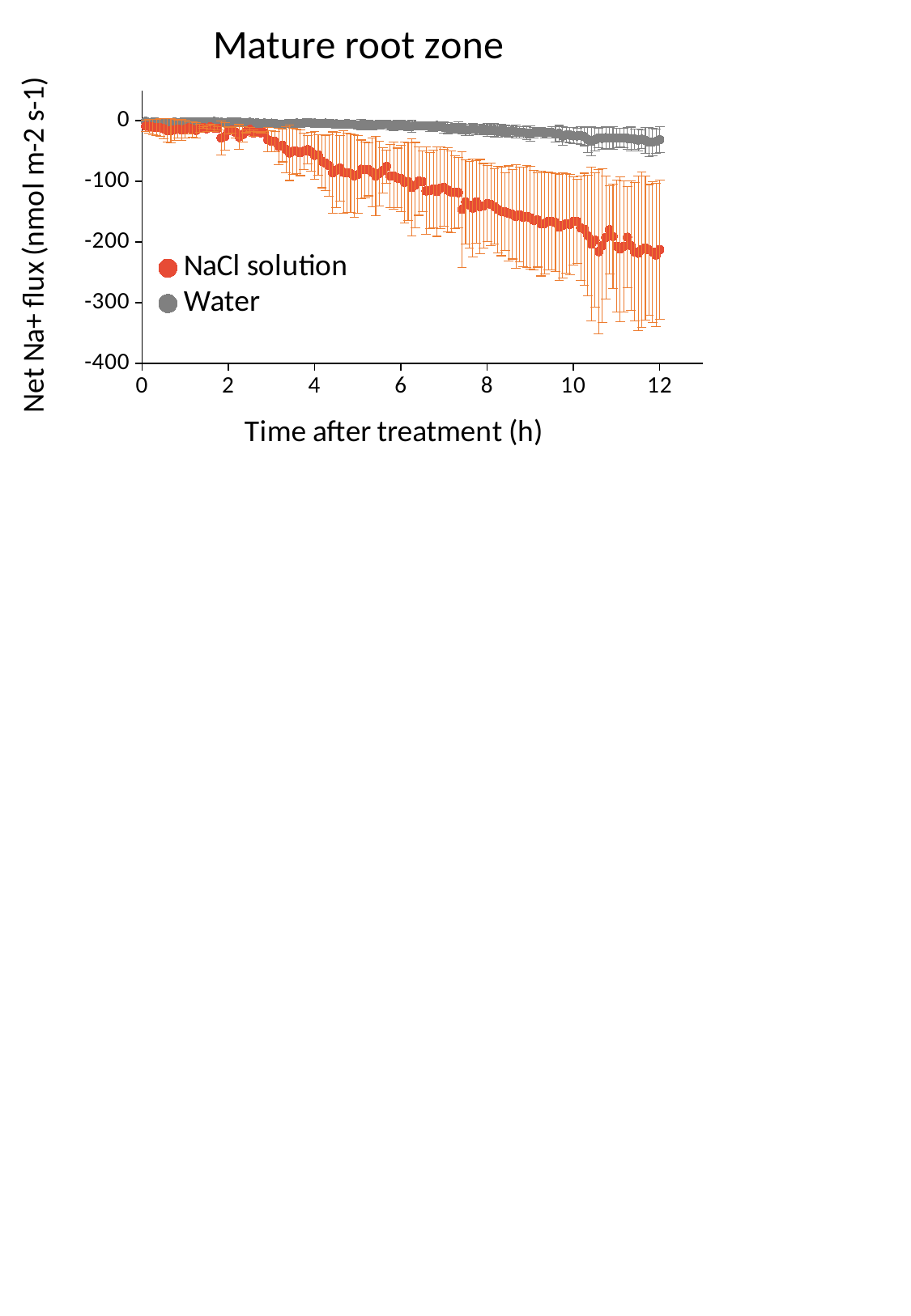

#### Chart: Mature root zone
| Category |
|---|

### Slide 2
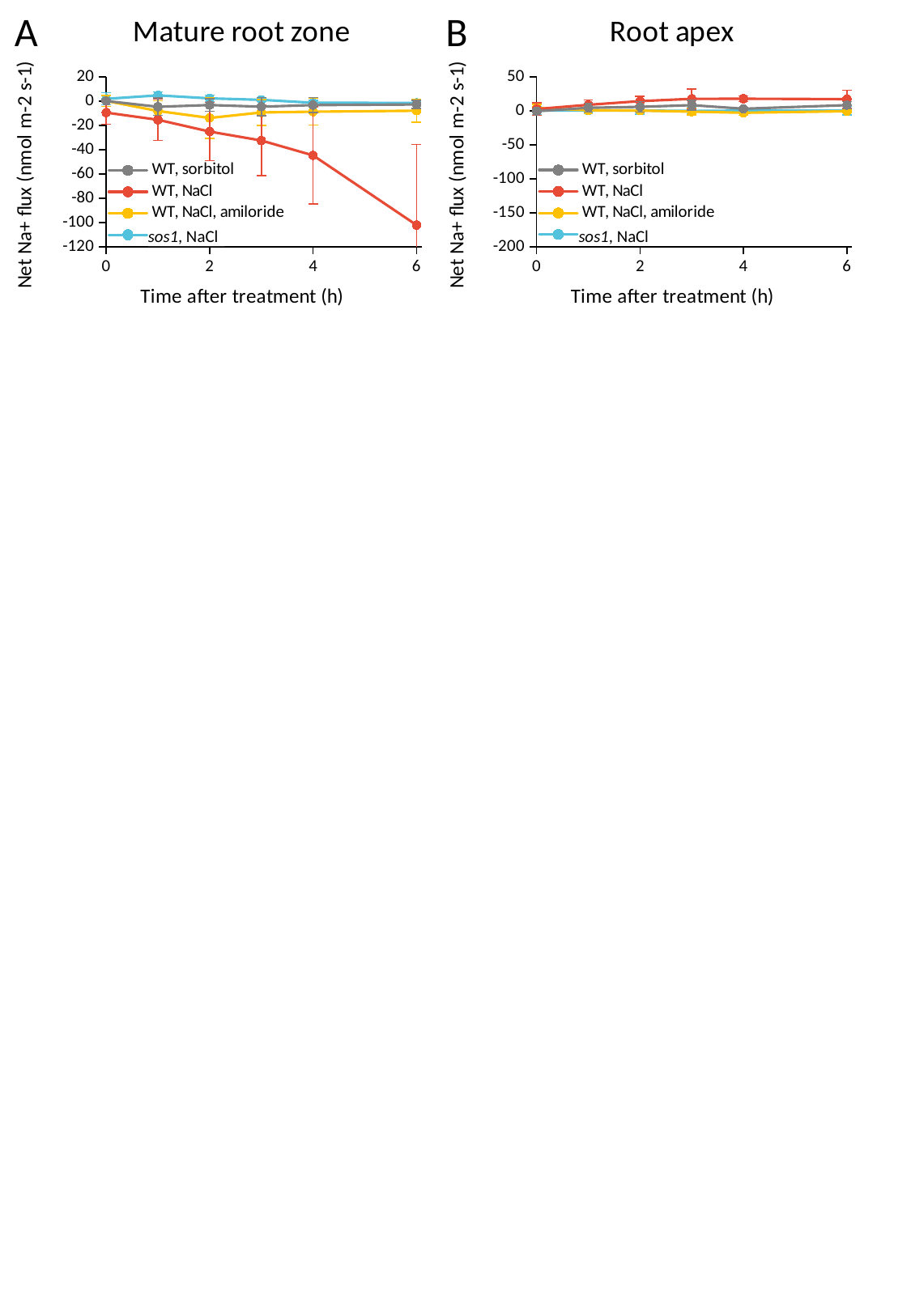

#### Chart: Mature root zone
| Category | | | | |
|---|---|---|---|---|A
#### Chart: Root apex
| Category | | | | |
|---|---|---|---|---|B
sos1, NaCl
sos1, NaCl

### Slide 3
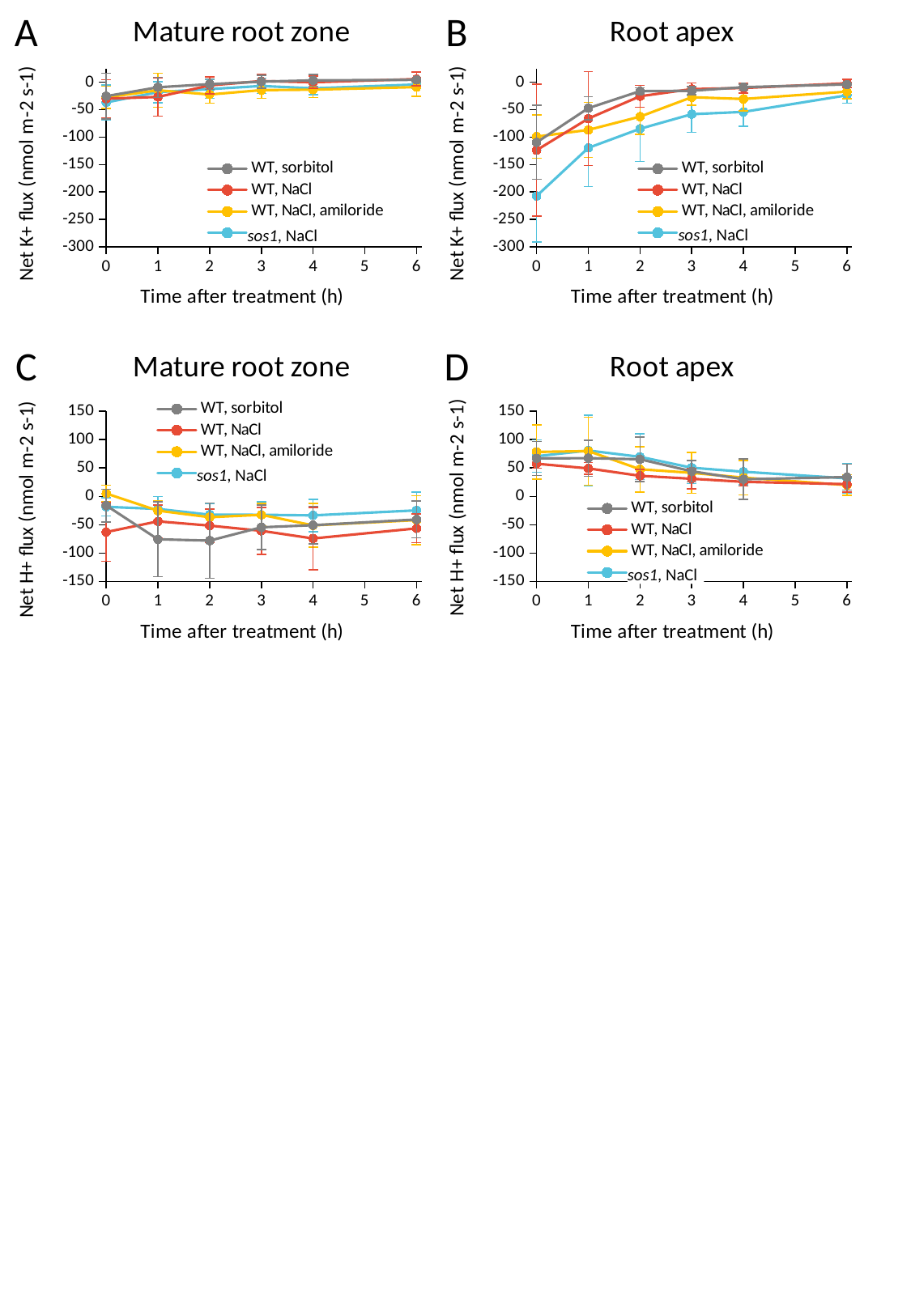

#### Chart: Mature root zone
| Category | | | | |
|---|---|---|---|---|A
#### Chart: Root apex
| Category | | | | |
|---|---|---|---|---|B
sos1, NaCl
sos1, NaCl
D
#### Chart: Root apex
| Category |
|---|
#### Chart: Mature root zone
| Category | | | | |
|---|---|---|---|---|C
sos1, NaCl
sos1, NaCl

### Slide 4
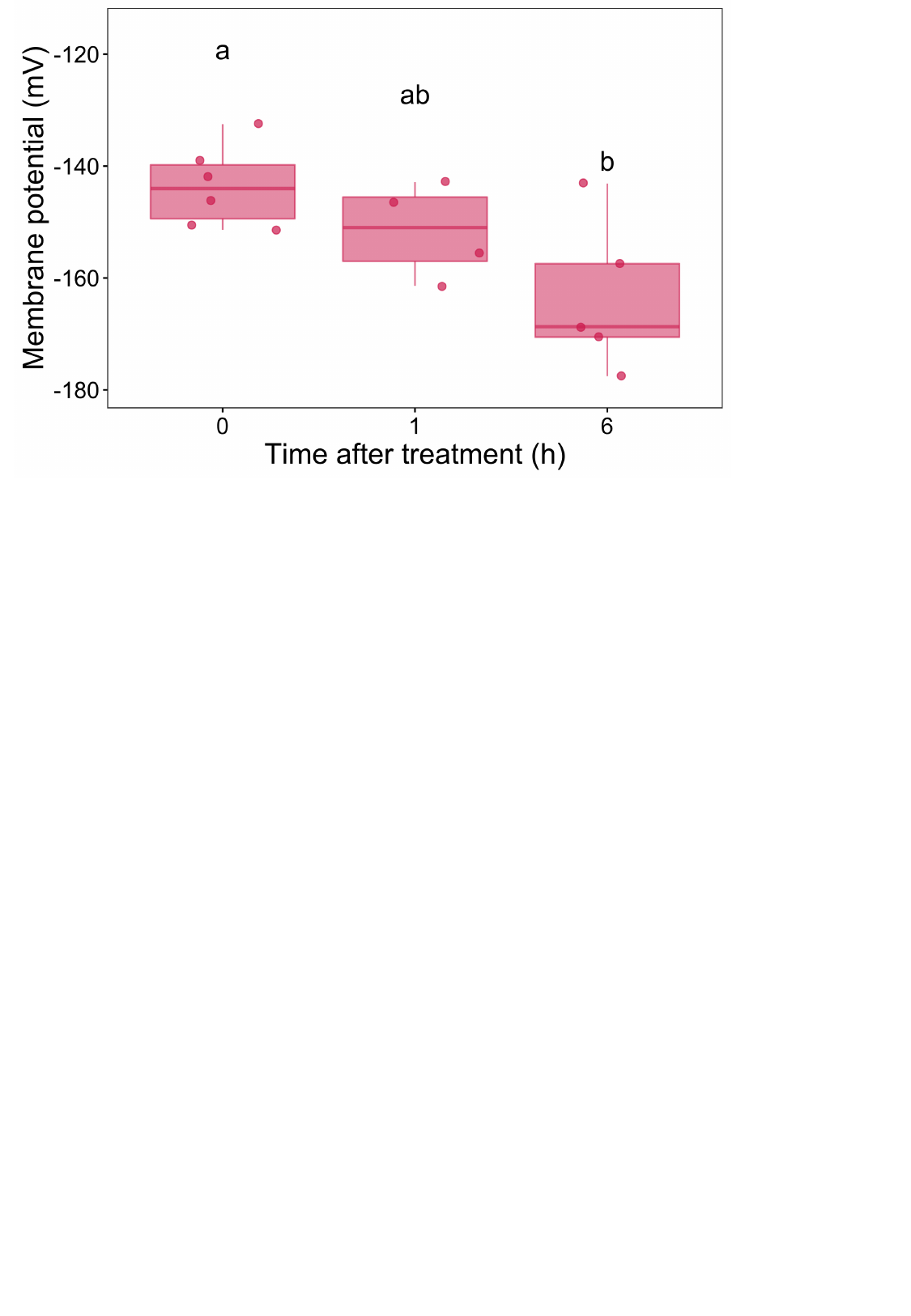
